# Ocular Motoneuron Pools Develop Along a Dorsoventral Axis in Zebrafish, *Danio rerio*

**DOI:** 10.1101/049296

**Authors:** Marie R. Greaney, Ann E. Privorotskiy, Kristen P. D’Elia, David Schoppik

**Affiliations:** Departments of Otolaryngology, Neuroscience & Physiology, and the Neuroscience Institute New York University Langone School of Medicine, New York, NY 10016

## Abstract

Both spatial and temporal cues determine the fate of immature neurons. A major challenge at the interface of developmental and systems neuroscience is to relate this spatiotemporal trajectory of maturation to circuit-level functional organization. This study examined the development of two ocular cranial motor nuclei (nIII and nIV), structures in which a motoneuron’s identity, or choice of muscle partner, defines its behavioral role. We used retro-orbital dye fills, in combination with fluorescent markers for motoneuron location and birth-date, to probe spatial and temporal organization of the oculomotor (nIII) and trochlear (nIV) nuclei in the larval zebrafish. We described a dorsoventral organization of the four nIII motoneuron pools, in which inferior and medial rectus motoneurons occupy dorsal nIII, while inferior oblique and superior rectus motoneurons occupy distinct divisions of ventral nIII. Dorsal nIII motoneurons are, moreover, born before motoneurons of ventral nIII and nIV. Order of neurogenesis can therefore account for the dorsoventral organization of nIII and may play a primary role in determining motoneuron identity. We propose that the temporal development of ocular motoneurons plays a key role in assembling a functional oculomotor circuit.

## Introduction

Spatiotemporally-regulated neurogenic processes underlie the production of diverse neuronal classes and subclasses including those of the vertebrate retina (Cepko, 2014), cortical projection neurons (Desai and McConnell, 2000) andinterneu-rons (Sousa and Fishell, 2010), and individually-identifiable cells within the lineage of a *Drosophila* neuroblast (Novotny et al., 2002). Trajectories of sensory neuron development were recently linked to their mature sensitivity profiles (Li et al., 2012). In the motor system, links between spatiotemporal organization and functional circuitry exist among interneuron subtypes involved in motor circuits (Lewis and Eisen, 2003). In the zebrafish spinal cord, interneuron birth order is related not only to dorsoventral topography but also to electrophysiological properties and recruitment order during progressively faster swimming behaviors (Fetcho and McLean, 2010). Spatiotemporal organization has likewise been observed for mammalian spinal cord interneuron populations with distinct functional roles (Tripodi et al., 2011). Similar evidence at the level of motoneuron organization is sparse, but would be welcome given the clear correspondence of motoneuron-to-muscle wiring and behavioral outputs. This study seeks to address the outstanding question of developmental organization of ocular motoneuron pools.

The oculomotor periphery consists of three cranial motor nuclei whose connectivity has been well characterized across vertebrate species, thanks to their participation in highly conserved goal-driven and reflexive behaviors (Büttner-Ennever, 2006): the oculomotor (nIII), trochlear (nIV), and abducens (nVI) nuclei, nIII comprises motoneuron populations innervating four of the six extraocular muscles and is the only one of these nuclei to house multiple subpopulations. Motoneurons targeting each extraocular muscle are organized into pools within nIII, the coherence of and overlap between pools varying between species (Evinger, 1988). Such clustering also exists among motoneurons in the spinal cord, which form pools in a cadherin-dependent fashion that innervate distinct muscles (Price et al., 2002; Jessell et al., 2011; Demireva et al., 2011). The development of nIII motoneuron identity is controversial. One proposal for how individual motoneurons come to acquire their identity *is post-hoc* self-identification following random extraocular muscle innervation (Glover, 2003), consistent with target-derived signals regulating motoneuron connectivity (Ladle et al., 2007) but not with a stereotyped order of sub-population birth (Shaw and Alley, 1981). Others propose that a caudal-to-rostral order of differentiation of both extraocular muscle and motoneuron groups allows each newly-born pair to match (Evinger, 1988), although this conflicts with the observed order of motor nucleus development (Altman and Bayer, 1981; Varela-Echavarria et al., 1996). We set out to elucidate the process of motoneuron fate determination in larval zebrafish.

A model vertebrate, the larval zebrafish has contributed much to our understanding of the organization and development of oculomotor circuitry and behavior (Miri et al., 2011; Ma et al., 2014). Recent work described the timecourse of axonal pathfinding and synaptogenesis between oculomotor neurons and their target extraocular muscles (Clark et al., 2013), but no characterization of either nIII motoneuron pool organization or its development yet exists. As part of a rich behavioral repertoire (Fero et al., 2010), 3-4 day old larval zebrafish begin to perform eye movements (Easter Jr. and Nicola, 1997) involving all six extraocular muscles: two for nasal/temporal horizontal saccades (Schoonheim et al., 2010), and the remaining four for the vertical and torsional vestibulo-ocular reflexes (Riley and Moorman, 2000; Mo et al., 2010; Bianco et al., 2012). We took advantage of the external development, transparency, and genetic tractability of the larval zebrafish to probe nIII development using transgenic reporters of motoneuron location (Higashijima et al., 2000) and methods to mark birthdate *in vivo* (Caron et al., 2008).

Here we found that larval zebrafish ocular motoneuron pools in nIII are organized along a dorsoventral axis, and that a similar axis defines their birth order. We first measured the distribution of ocular motoneurons across nIII and nIV labeled in Tg(isl1:GFP) larvae, and used this data to construct a spatial framework for motoneuron localization. Using targeted retro-orbital dye fills, we localized ocular motoneuron somata in nIII and nIV of larval zebrafish. We found that nIII is divided into dorsal and ventral halves; nIV aligns with the dorsal half of nIII. Dorsal nIII contains mostly inferior/medial rectus (IR/MR) motoneurons, while inferior oblique (IO) and superior rectus (SR) motoneurons occupy largely distinct regions of ventral nIII. Next, we measured the time of terminal differentiation of motoneurons across nIII and nIV, and discovered a dorsoventral order to birthdate, complementary to the spatial organization of motoneuron pools. We propose that birth order is the primary determinant of the spatial organization and fate of ocular motoneurons in nIII.

## Methods

### Fish husbandry

All protocols and procedures involving zebrafish were approved by the New York University Langone School of Medicine Institutional Animal Care & Use Committee (IACUC). All larvae were raised at 28.5°C at a density of 20-50 larvae in 25-40mL of buffered E3 (1mM HEPES added). Older larvae, >7 days post-fertilization (dpf), were raised in the facility on a standard 14/10 hour light/dark cycle, and fed twice daily.

### Transgenic lines

Experiments were done on the mitfa-/- background to remove pigment. Larvae homozygous for Tg(isl1:GFP) (Higashijima et al., 2000) were used to label cranial motoneurons for retro-orbital fill. Homozygotes were identified by level of GFP fluorescence relative to heterozygous siblings. Larvae for photoconversion experiments were doubly-heterozygous for Tg(huC:Kaede), (Sato et al., 2006) which labels post-mitotic neurons, and Tg(isl1:GFP).

### Retro-orbital fills

Larvae were anesthetized in 0.02% Ethyl-3-aminobenzoic acid ethyl ester (MESAB, Sigma-Aldrich E10521) at 4-5 dpf or 13 dpf until no longer responsive to touch/dish taps. Anesthetized larvae were mounted in 2% low-melting point agar (ThermoFisher Scientific 16520), positioned with the right eye up and oriented for best access to nerve targeted (Figure 1A). Larvae were oriented predominantly dorsal up for superior oblique (SO) and superior rectus (SR) neuron fills, or predominantly ventral up for inferior oblique (IO) and inferior/medial rectus (IR/MR) fills, tilted slightly off-axis in either case. Agar was cleared from the area above and around the muscle innervated by the nerve of interest. An electrochemically-sharpened tungsten needle (10130-05, Fine Science Tools) was used to create an incision in the skin at the position of the muscle, closely following the outside of the eye (Figure 1A–1B). Any excess liquid was removed from the area. A second sharpened tungsten needle was used to hold a piece of solidified dextran-conjugated Alexa Fluor 647 dye (10,000 MW, ThermoFisher Scientific D-22914) into the incision, until the dye had spread such that the incision location was fully colored (Figure 1A”). Larvae were left for a minimum of five minutes following dye application, at which point E3 was added over the agar drop. Larvae were then freed from the agar and allowed to recover in E3 for a minimum of four hours, and usually overnight.

**Figure 1:**
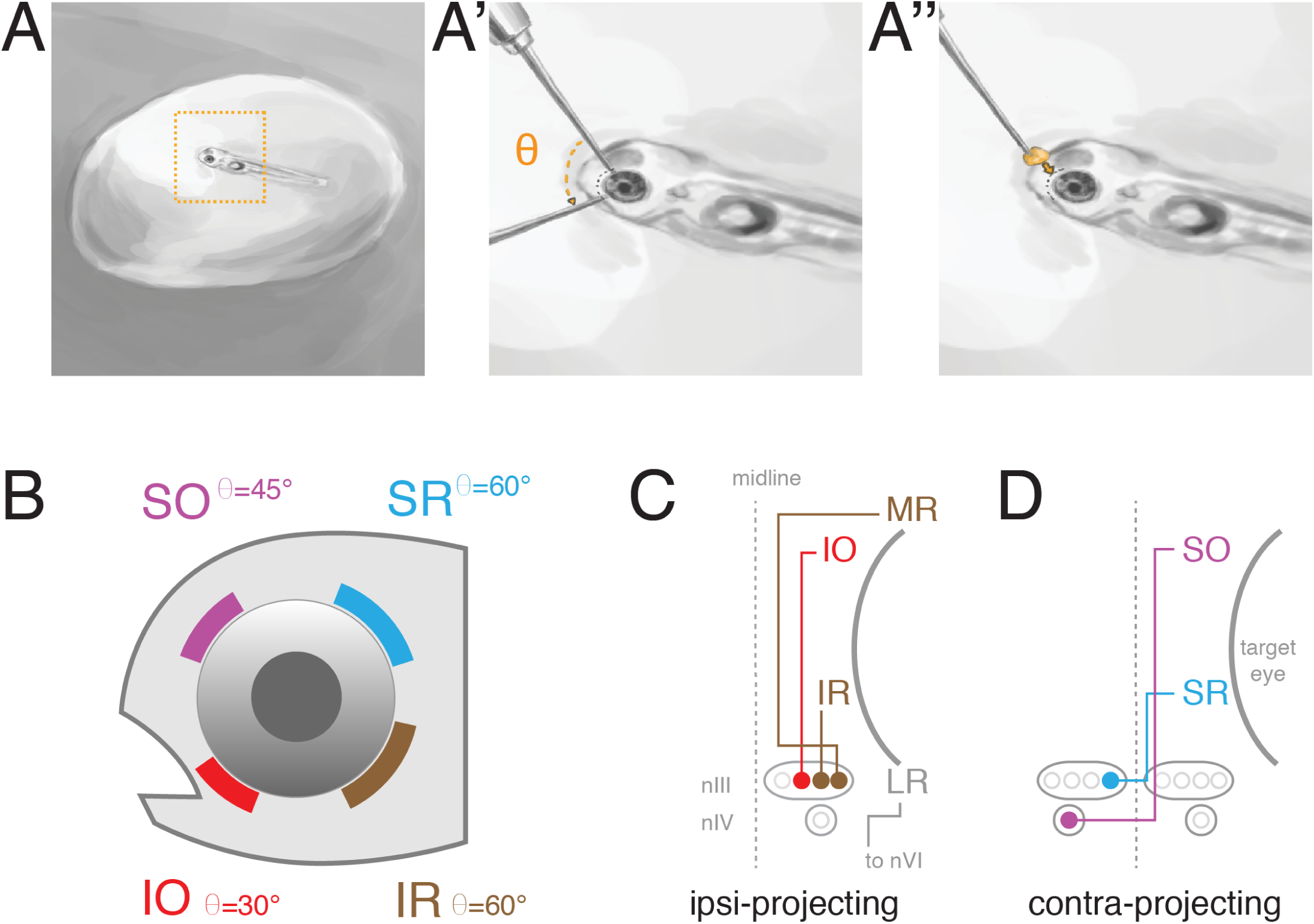
Using retro-orbital dye fills to label ocular motoneurons. A) Retro-orbital fill procedure. Larvae were anesthetized and immobilized in 2% agar, target eye near the surface (A). Agar was cleared from eye and a tungsten needle used to make an incision of angle (Θ) in the vicinity of one/more extraocular muscle (A’). Crystallized dye was placed at incision site (A”); somata were imaged later. B) Targeted position and angle (Θ) of incision around eye to label a given motoneuron population. C) Schematic of extraocular muscle innervation by ipsilaterally-projecting motoneurons. LR motoneurons located in nVI were not targeted for dye fills. D) Schematic of extraocular muscle innervation by contralaterally-projecting motoneurons. MR IR SR LR medial, inferior, superior, lateral rectus. SO, IO: superior, inferior oblique, nIII: oculomotor nucleus. nIV: trochlear nucleus. nVI: abducens nucleus.

### Data collection and analysis

#### Assignation of filled cells as specific motoneurons

Larvae were anesthetized in 0.02% MESAB and mounted dorsally in 2% agar for image collection. Larvae were imaged between 5-7 dpf, or at 14 dpf, using the following confocal microscopes: Zeiss LSM 510 (SO and SRdata, 40x/0.8NA objective), Leica SP8 in photon counting mode (IO data, 20x/1.0NA objective), Zeiss LSM 710 (IR/MR data, 40x/1.1NA objective), and Zeiss LSM 800 (Kaede data, 40x/1.0NA objective). Image analysis was performed using ImageJ (Fiji) (Schindelin et al., 2012) and cell location recorded on vector templates using Illustrator CC 2014 (Adobe Systems Inc, San Jose, CA).

To identify particular somata as motoneurons associated with a given ocular muscle, we used two criteria: first, the projection patterns of ocular motoneurons (Figure 1C–1D), which are highly conserved across the vertebrate kingdom (Evinger, 1988); and second, the known lack of GFP expression in IO motoneurons in the Tg(isl1:GFP) background (Higashijima et al., 2000). Somata filled contralaterally to the right eye (left side of the brain) were defined as SO motoneurons if found in nIV, or SR motoneurons if found in nIII (Figure 2A). Somata filled ipsilaterally to the right eye (right side of the brain) were defined as IR/MRmotoneurons, if GFP+, or IO motoneurons, if GFP-(Figure 2B). We observed 3 putative contralateral IO motoneurons, which were excluded.

**Figure 2:**
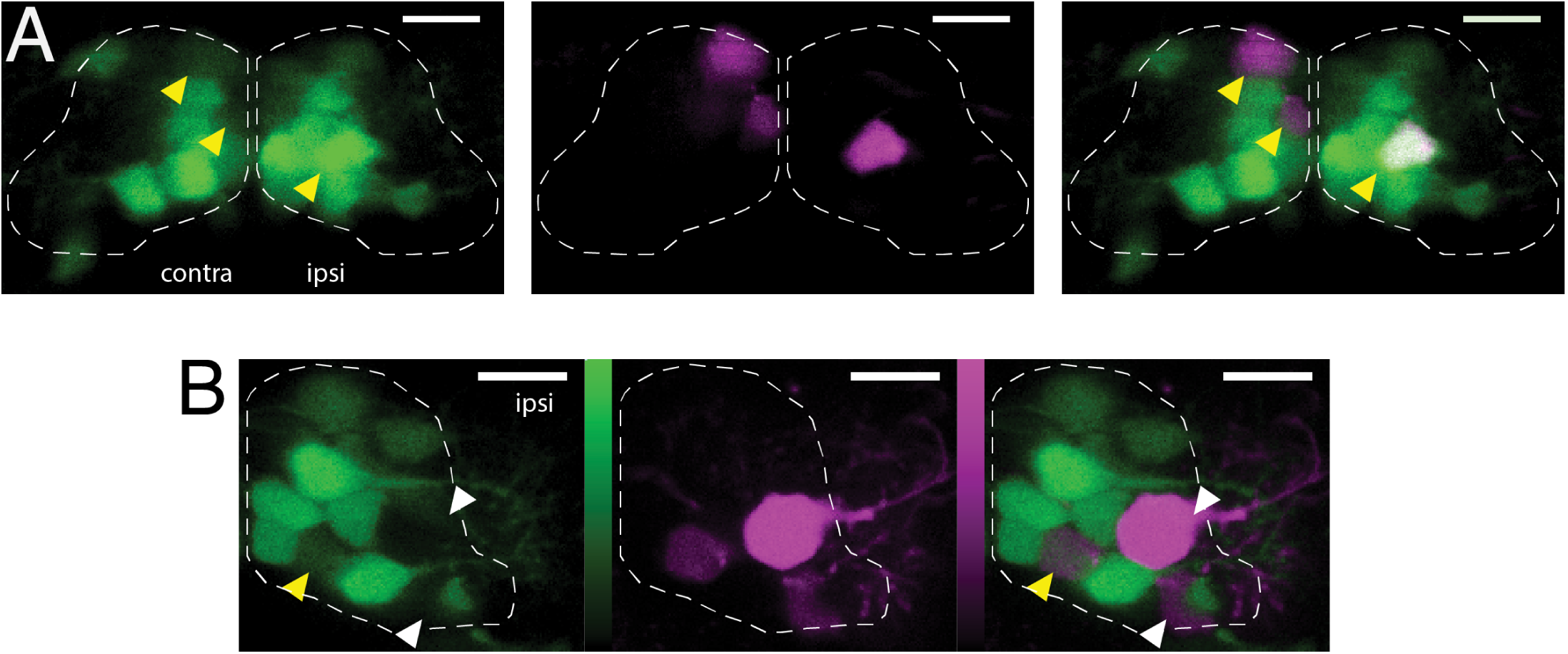
Identification of dye-filled motoneuron subtypes by projection pattern and Tg(isl1:GFP) expression. A) Filled motoneurons in ventral nIII (plane z4, dashed line). Yellow arrowheads indicate dye-filled motoneurons that were also GFP+ (contralateral: SR motoneurons; ipsilateral: IR/MR motoneurons). B) Filled motoneurons in ventral nIII (plane z5, dashed line). White arrowheads indicate dye-filled motoneurons without GFP (IO motoneurons); yellow arrowhead indicates a dye-filled motoneuron that was also GFP+ (IR/MR motoneuron). Color bars represent photons detected in photon-counting mode (IO data only); green range: 0-198; magenta range: 0-231. Green = GFP; magenta = Alexa Fluor 647 dye. Scale bars: 10 *μm*.

#### nIII/nIV template generation and motoneuron identification

To provide a consistent spatial framework for placement of filled cells within the boundaries of nIII/nIV, we generated a template (Figure 3). Two evenly aligned stacks, showing minimal roll or pitch deviations, were chosen to estimate stereotyped boundaries of GFP+ cells. 15 dorsoventral divisions of the nuclei were established, spaced approximately 6 *μm* apart (“z0-z14”). Maximum intensity projections were made using 12 *μm* of data centered at each dorsoventral division. Boundaries were manually drawn around the extent of GFP+ cells within each projection and expanded to encompass all cells across the two larvae to represent the expected extent of GFP+ cells at each division. The midbrain-hindbrain boundary delineated nIII and nIV. To test how stereotyped GFP+ expression relative to anatomical landmarks was across larvae, we manually bounded both green somatic fluorescence and the ascending posterior mesencephalic central artery (PMCtA) on each side of the midline, at the level of the medial longitudinal fasciculus (MLF), and measured the distance between the center of fluorescence intensity and the centroid of the corresponding PMCtA (Figure 3B).

**Figure 3:**
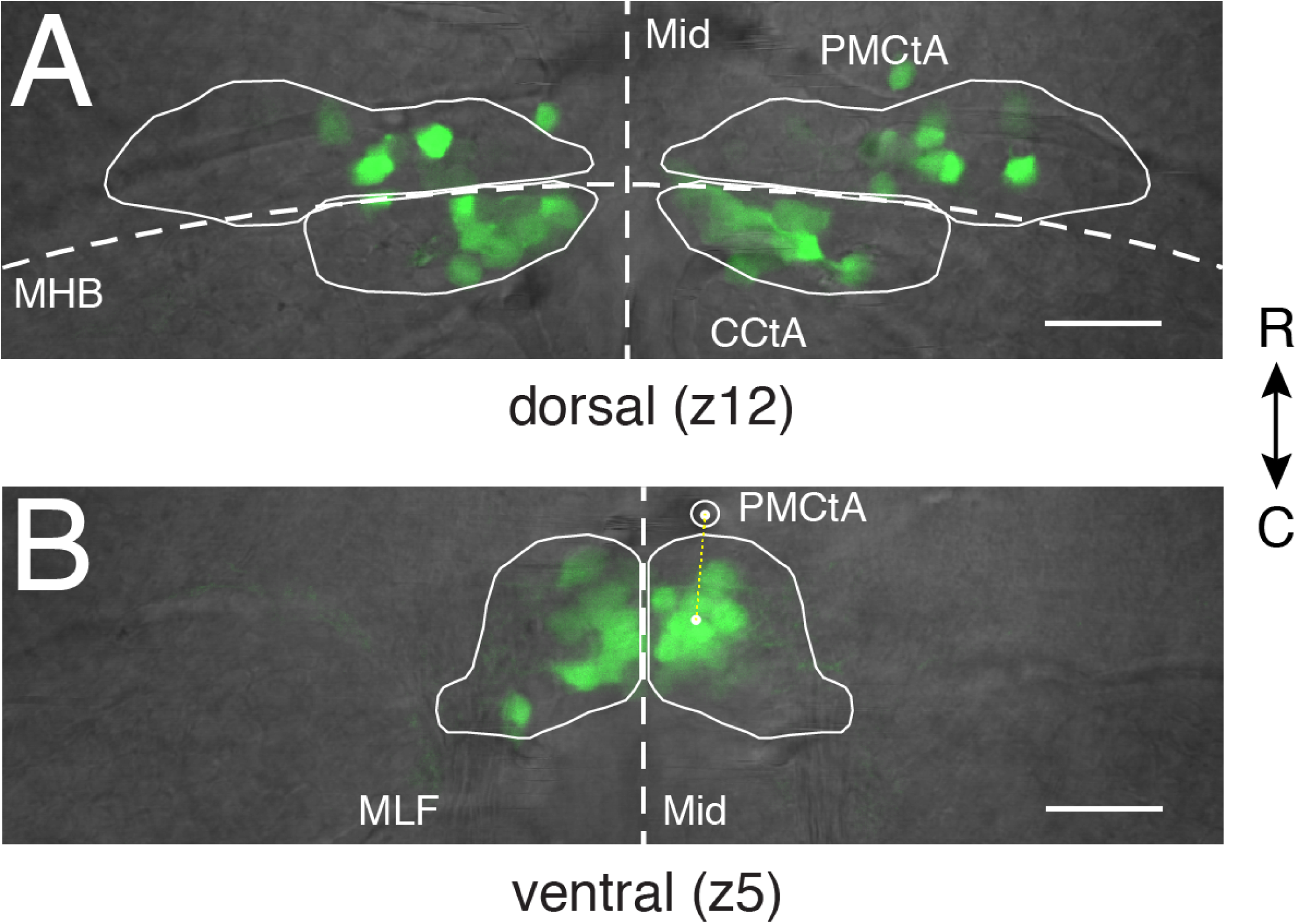
Fluorescent somata in Tg(isl1:GFP) zebrafish define the spatial extent of ocular motoneurons in nIII and nIV. A) Positions of fluorescent motoneurons in dorsal nIII/nIV (z12) relative to anatomical landmarks. B) Positions of fluorescent motoneurons in ventral nIII (z5) relative to anatomical landmarks. Yellow dotted line represents the distance between the center of intensity of green fluorescence right of the midline and the centroid of the right PMCtA. Mid: midline. MHB: midbrain-hindbrain boundary. MLF: medial longitudinal fasciculus. PMCtA: posterior mesencephalic central artery. CQA: cerebellar central artery. Scale bars: 20 *μm*.

When excited at 488 nm, Alexa Fluor 647 dye emits fluorescence detectable in the 500-550 nm collection channel at the gains necessary to resolve dim signal in Tg(isl1:GFP) larvae. We developed a thresholding procedure to differentiate GFP emission from dye emission. Control fills were first performed and imaged in Tg(isl1:GFP) negative siblings. Pixel values from brightly fluorescent dye-filled cells were used to establish an intensity floor for the GFP channel. For each data stack, imaged under identical experimental conditions, the intensity was adjusted to only display values above the floor. Applying this minimum intensity display threshold to stacks from homozygous Tg(isl1:GFP) larvae allowed us to distinguish dye-filled GFP-cells from faintly GFP+ cells. To define somatic location within nIII/nIV, individual filled cells were classified as particular motoneurons as described above and localized within the boundaries of the template at the appropriate dorsoventral plane. Somata were represented as circles approximately 4 *μm* in diameter, roughly the size of a small nIII/nIV neuron. Neurons with somata that sat between template planes were assigned based on the plane of brightest fluorescence. Each soma was only counted once.

#### Birthdating of neurons using a photolabile protein

Larvae heterozygous for Tg(isl1:GFP) and Tg(huC:Kaede) were exposed for 10 minutes to focused full spectrum light from a mercury light source at 22, 29, 36, 43, or 50 hpf (Figure 4A) to photoconvert green Kaede to red. Larvae were kept in the dark following collection until anesthesia and mounting for imaging at 5 dpf. As per the Birthdating Analysis by photocon-verted fluorescent Protein Tracing In vivo, with Subpopulation Markers (BAPTISM) procedure, larvae were imaged in both the red and green channel twice in succession at 5 dpf (Caron et al., 2008). The first image stack contained red Kaede emission only in neurons that were already post-mitotic during the earlier photoconversion, and green Kaede emission from both unconverted neurons and cranial motoneurons (GFP) (4B, left column). Next, all remaining unconverted Kaede was photo-converted from green to red using a 405 nm laser. Lastly, a second stack was taken with the same settings to obtain data comparable to the first stack (4B, right column). There, the only remaining green was from Tg(isl1:GFP) neurons. Comparison of the red channel from the first stack (converted Kaede) and the green channel from the second stack (GFP) permitted identification of cranial motoneurons that were post-mitotic by the time of initial photoconversion.

**Figure 4:**
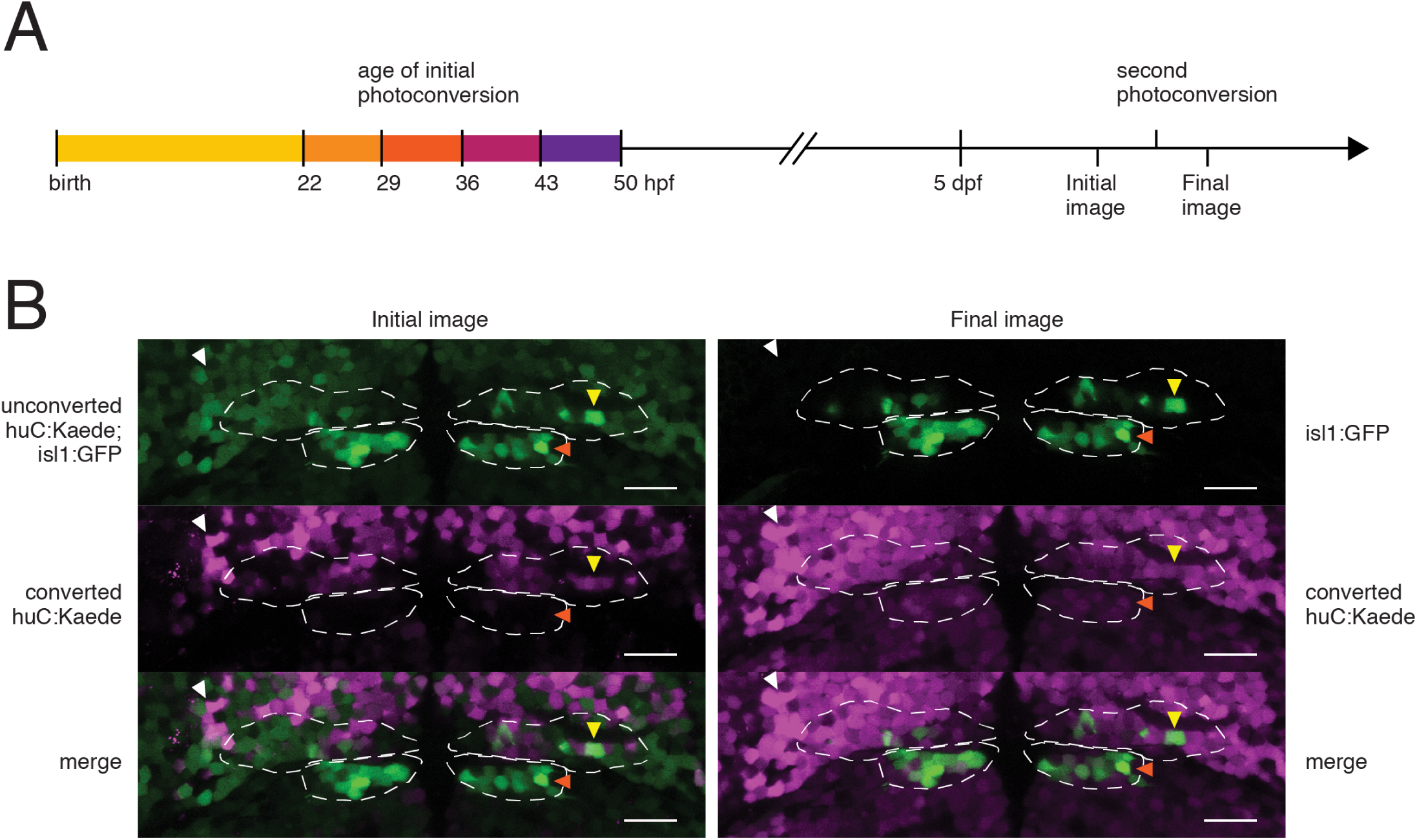
Birthdating Analysis by photoconverted fluorescent Protein Tracing In vivo, with Subpopulation Markers (BAPTISM) method (Caron et al., 2008) for identifying the time of terminal differentiation (birthdate) of motoneurons in Tg(huC:Kaede; isl1:GFP) zebrafish, which express photolabile fluorescent protein Kaede pan-neuronally. A) Neurons born by initial photoconversion (one of five time points shown) contain converted Kaede in Initial image, taken at 5 dpf. Second photoconversion converts remaining Kaede, leaving GFP as the only green signal in Final image. Images are compared to determine birthdate of GFP+ motoneurons. B) Dorsal plane (z12, dashed line) from a 5 dpf Tg(huC:Kaede; isl1:GFP) larva initially photoconverted at 36 hpf. Yellow arrowheads indicate an nIII motoneuron (isl:GFP+, Final image) born by 36 hpf (converted huC:Kaede+, Initial image). White arrowheads indicate a neuron born by 36 hpf not belonging to nIII/nIV. Orange arrowheads indicate an nIV motoneuron not born by 36 hpf. Scale bars: 20 *μm*.

To establish a threshold for red Kaede signal that corresponded to a post-mitotic neuron, rather than basal photoconversion, we used data from stacks of Tg(huC:Kaede);Tg(isl1:GFP) siblings that had not been exposed to light. Control and experimental stacks were matched for overall Kaede expression. 7 experimental stacks that greatly exceeded control stacks in overall Kaede intensity were not used for analysis. GFP+ neurons in the final image were placed into the map of nIII/nIV only if they contained supra-threshold red Kaede signal from the initial image.

## Results

### The dorsoventral distribution of isl1:GFP+ cells in cranial nuclei III and IV

To establish a spatial framework to localize ocular motoneurons, we employed the Tg(isl1:GFP) line, which labels SO, SR, IR, and MR motoneurons (Higashijima et al., 2000). Ocular motoneurons were observed 100-200 *μm* deep in the midbrain and hindbrain. We divided a region that spanned about 90 *μm* in depth into 15 unique planes spaced every 6 *μm*, and defined bounds around GFP+ motoneurons (Figure 3). We tested the consistency of GFP+ expression relative to other anatomical landmarks across larvae by measuring the distance between the center of intensity of green somatic fluorescence on one side of the midline and the center of the corresponding ascending PMCtA at the level of the MLF (Figure 3B). We measured this distance for each hemisphere in 10 larvae and recorded an average of 18±4 *μm* (SD). We conclude that the distribution of GFP fluorescence is largely stereotyped relative to anatomical landmarks identifiable under visible illumination.

To quantify motoneurons in nIII and nIV, we characterized the distribution of GFP+ motoneurons within nIII and nIV in ten 5 dpf larvae (Figure 5). We observed 104±20 (SD) nIII motoneurons (max: 134, min: 68) and 47±8 nIV motoneurons (max: 58, min: 35) per side per larva. Note that nIII totals do not include IO motoneurons, which are not labeled in Tg(isl1:GFP) (Higashijima et al., 2000). GFP+ neurons were found in comparable numbers across larvae at every plane (Figure 5B). As in avians and other teleosts, GFP+ motoneurons in nIII could be organized into dorsal and ventral sections (Evinger, 1988). Here, each division accounts for approximately half of the height and cell count of the nucleus. GFP+ motoneurons in nIV align with the dorsal half of nIII on the caudal side of the midbrain-hindbrain boundary (MHB), and number just under half as many as nIII.

**Figure 5:**
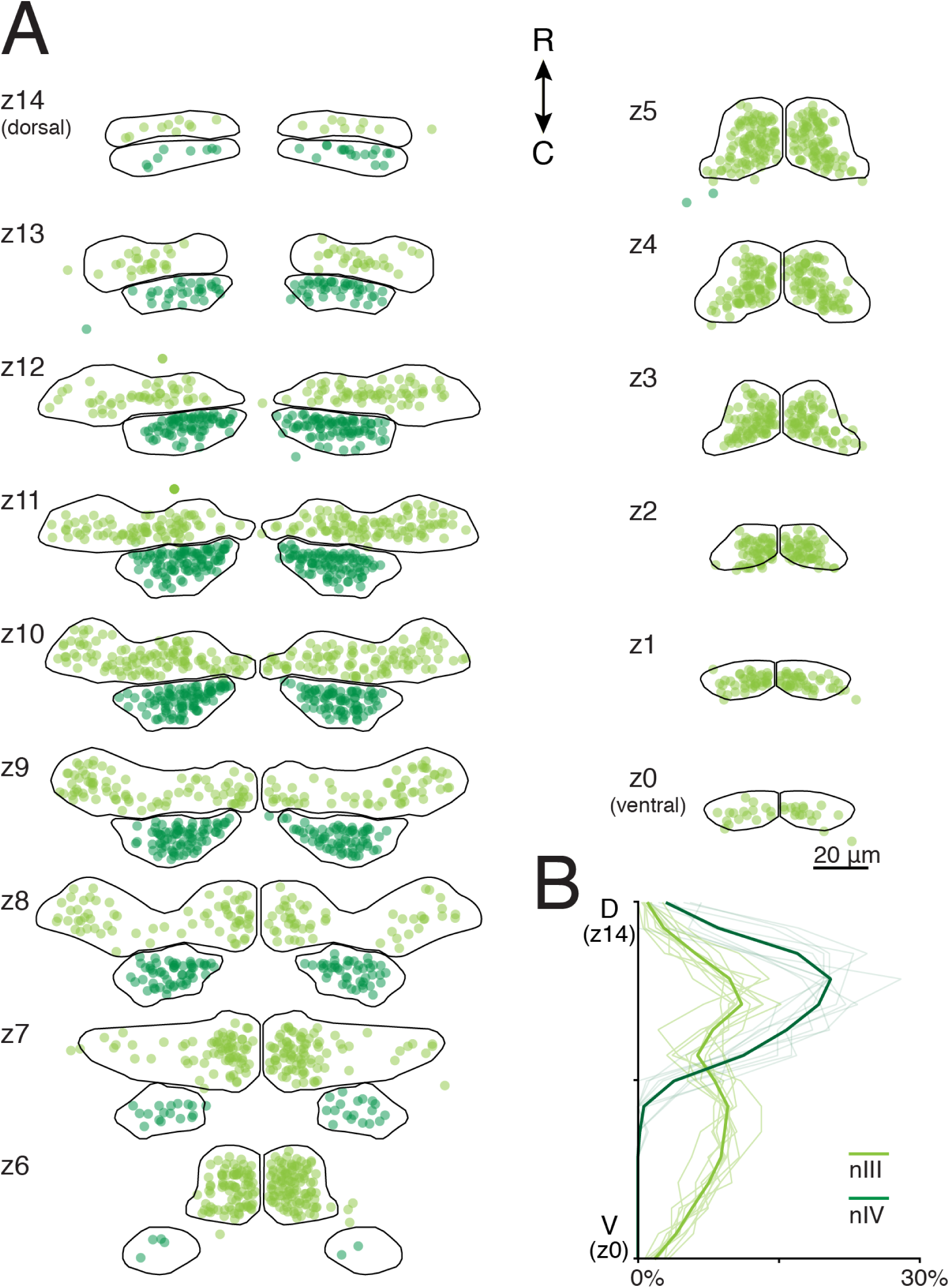
Distribution of labeled motoneuron somata in 5-7 day old Tg(isl1:GFP) zebrafish. A) GFP+ motoneuron somata are shown as circles in nIII (light green) and nIV (dark green). The dorsoventral extent of nIII/nIV is subdivided into 15 6-*μm*-thick planes, labeled z14 (most dorsal) - z0 (most ventral). B) Mean and individual (background traces) probability distributions from labeled motoneurons across nIII/nIV. nIII: n = 10 larvae, 2080 cells. nIV: n = 10 larvae, 936 cells.

As the level of GFP varies across motoneurons, we tested the inter-observer reliability. Motoneurons from the same five larvae were counted independently by two observers (MG and AP). A 3-way analysis of variance grouping data for dorsoventral sublevel, larva, and observer rejected the null hypothesis of equal means for all three variables (*p* < 10^−16^, *p* < 10^−16^, *p* = 0.044, *df* = 14, 4, 1, *F* = 41, 16, 4.1). Since we cannot rule out an observer effect on motoneuron quantification, we measured inter-observer differences. On average, MG found 9% more GFP+ cells, but inter-observer counts were correlated with a pairwise linear correlation coefficient of 0.97. We conclude that while the absolute motoneuron counts may vary by experimenter, the relative spatial distribution of motoneurons is robust to inter-experimenter variation. Bounds representing the extent of isl1:GFP+ motoneurons are a suitable spatial framework to map motoneuron position.

### Superior rectus motoneurons are restricted to ventral nIII in both 5-7 and 14 day-old zebraflsh

To determine if Tg(isl1:GFP) is a veridical marker of ocular motoneurons, we targeted SO motoneurons for retro-orbital fills. SO motoneurons comprise the entirety of nIV and project to the contralateral eye with distinct dorsal axonal morphology. Our fills labeled individual somata within a cluster just caudal to the MHB and contralateral to the targeted eye. Each dye-filled soma was also GFP+ in this area (Figure 6, magenta). Dye-filled axons were often visible within the GFP+ IVth nerve, ascending dorsally and contralaterally We concluded from their GFP expression, axonal morphology and location in the hindbrain that these cells were successfully-filled SO motoneurons. We never observed filled somata in nIV that were not GFP+, supporting the hypothesis that the Tg(isl1:GFP) is a veridical marker of motoneurons. On average, we labeled 42±33 motoneurons per larva, 14±10 of which were contralateral neurons in nIV, yielding a total of 606 SO motoneurons across 42 larvae. We observed a maximum of 43 SO motoneurons in a single larva. We conclude that the retro-orbital fill method is well-suited to stochastically label ocular motoneurons.

**Figure 6:**
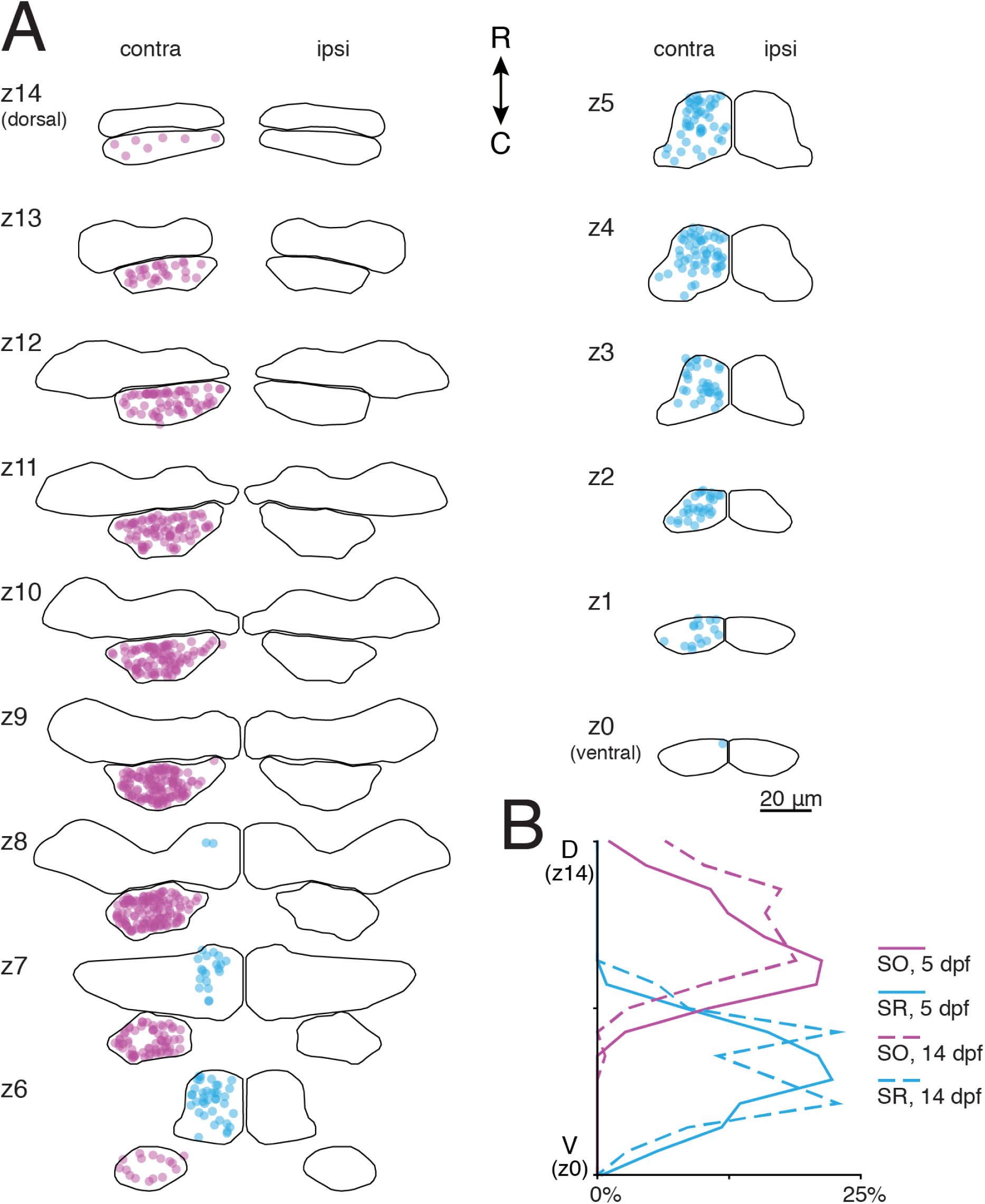
Superior rectus motoneurons are located exclusively in ventral nIII in both 5-7 and 14 day old zebrafish; superior oblique motoneurons are located in nIV. A) Location of SO (magenta) and SR (blue) motoneurons at 5-7 dpf across nIII/nIV. GFP+ neurons in nIII contralateral to the filled eye were defined as SR motoneurons; GFP+ neurons in nIV contralateral to the filled eye were defined as SO motoneurons. B) Probability distributions of dye-filled SO and SR motoneurons across nIII/nIV at 5-7 dpf (solid lines) and 14 dpf (dashed lines). SO motoneurons: n = 42 larvae, 606 cells (5-7 dpf); n = 19 larvae, 138 cells (14 dpf). SR motoneurons: n = 22 larvae, 229 cells (5-7 dpf); n = 14 larvae, 35 cells (14 dpf).

To examine nIII subpopulation distributions, we first defined SR-projecting motoneurons as all GFP+ nIII motoneurons filled on the contralateral side of the brain. SR motoneurons appeared solely in the ventral half of nIII and were particularly dense in medial and rostral portions of the nucleus (Figure 6, cyan). On average, we labeled 60±44 neurons per larva, 10±12 of which were contralateral GFP+ neurons in nIII, yielding a total of 229 SR motoneurons across 22 larvae. We observed a maximum of 41 SR motoneurons in a single larva; both dorsolateral and within-subdivision distribution persisted if this larva was removed from consideration. We did not observe any GFP-filled cells in this part of contralateral nIII, supporting the hypothesis that, as with SO motoneurons, Tg(isl1:GFP) reliably labels SR motoneurons.

Early in development, the ocular motoneuron somata may still be motile (Puelles-Lopez et al., 1975). To determine if the spatial organization at 5-7 dpf represented stable localization of motoneurons within the nucleus, we filled SO and SR motoneurons in 13 dpf larvae. Contralaterally-filled cells were distributed with similar frequency among dorsoventral subdivisions at 14 dpf as at 5 dpf (Figure 6) in similar positions within subdivisions, supporting the hypothesis that the organization evident among motoneurons at 5-7 dpf is stable across early development.

### Inferior oblique motoneurons are found in ventral nIII, largely distinct from superior rectus motoneurons, while inferior/medial rectus motoneurons are found in dorsal nIII

We next defined IO-projecting motoneurons as all GFP-somata filled on the ipsilateral side of the brain. Similar to SR motoneurons, the large majority of IO neurons lay in the ventral half of nIII at 5-7 dpf. A few scattered IO cells were also present in dorsal nIII, interspersed within the GFP+ cell population. Most of the ventral IO motoneurons were located on the lateral or caudal edges of the GFP+ cell cluster, although some were found closer to the midline (Figure 7A, red). 154/328 labeled neurons were GFP-in 30 larvae; all but one of the larvae had GFP+ neurons labeled as well. On average, we labeled 11±7 neurons per larva, 5±4 of which were GFP-neurons in nIII. The maximum number of IO neurons filled in a single larva was 13. The low number is likely due to the conservative criteria used to define a soma as GFP-, given the presence of GFP+ neuropil and the axial resolution of the confocal microscope.

**Figure 7:**
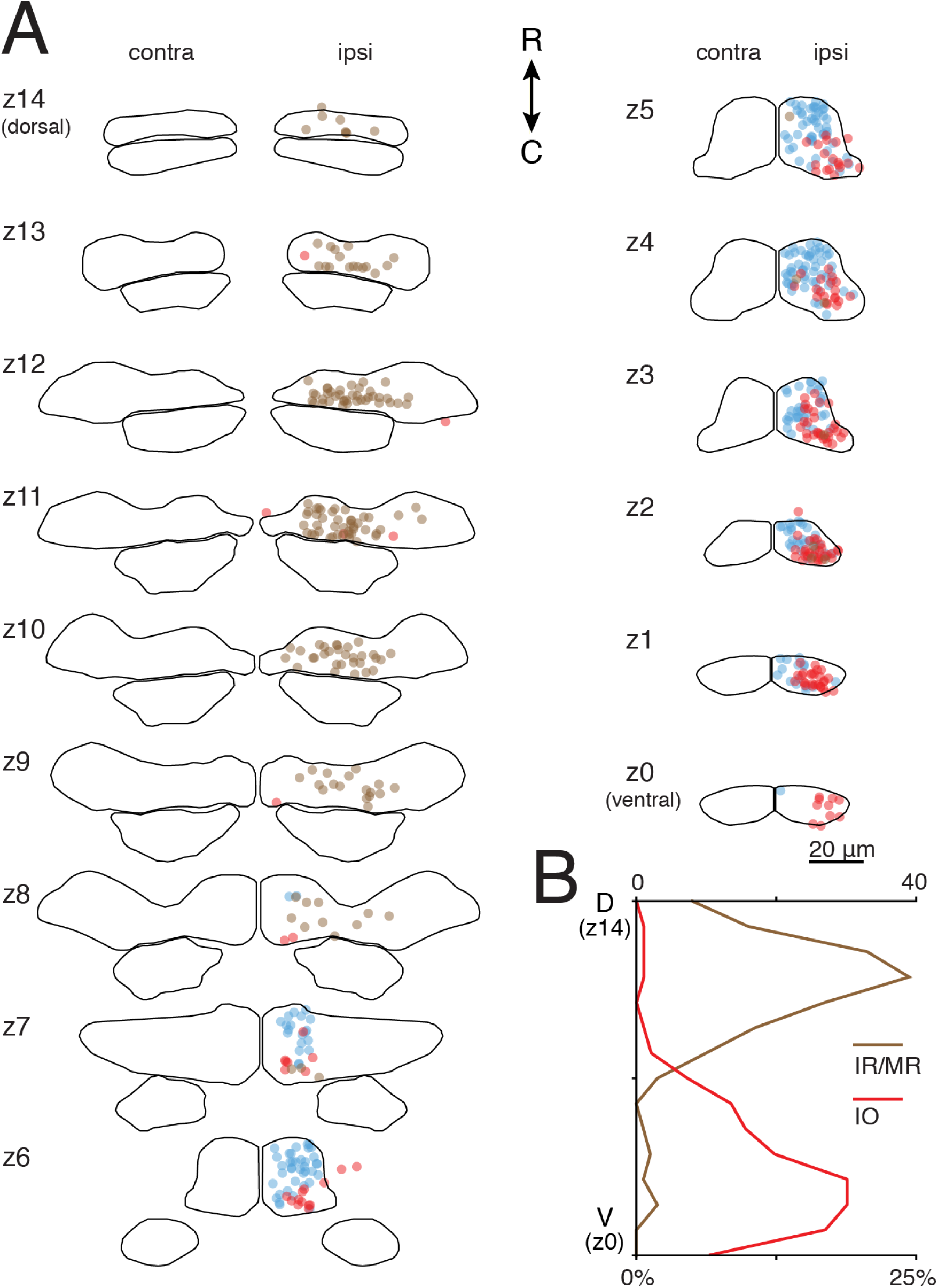
Inferior oblique motoneurons are located mainly in ventral nIII in 5-7 day old zebrafish, in a distinct caudal region relative to superioi rectus motoneurons, while inferior/medial rectus motoneurons are located mainly in dorsal nIII. A) Location of IO (red), SR (blue), and IR/MR (brown) motoneurons at 5-7 dpf across nIII/nIV. GFP-neurons in nIII ipsilateral to the filled eye were defined as IO motoneurons: GFP+ neurons in nIII ipsilateral to the filled eye were defined as IR/MR motoneurons. SR data is mirrored across the midline from Figure 4. B) Probability distributions of dye-filled IO and IR/MR motoneurons across nIII/nIV. IO motoneurons: n = 30 larvae (5-7 dpf), 154 cells. IR/MR motoneurons: n= 17 larvae (5-7 dpf), 160 cells.

Data from adult teleosts suggests that IO and SR motoneuron pools are spatially distinct (Graf and McGurk, 1985). We compared the positions of IO and SR motoneurons within ventral nIII subdivisions (Figure 7). As in adult teleosts, we observed that IO and SR populations are largely spatially segregated into caudolateral and rostromedial divisions of ventral nIII, respectively.

The remaining nIII subpopulations, IR and MR motoneurons, both innervate ipsilateral extraocular muscles, and are both labeled in Tg(isl1:GFP). Furthermore, both the IR and MR muscles are found near the dye insertion site (Kasprick et al., 2011). Thus, we were unable to distinguish IR from MR motoneurons in our fills. To optimally differentiate IR/MR motoneurons from IO motoneurons, we only evaluated larvae that had no dye in GFP-somata. IR/MR motoneurons were located throughout the entirety of dorsal nIII at 5-7 dpf, and infrequently in ventral nIII (Figure 7). On average, we labeled 10±11 neurons per larva, 9±11 of which were ipsilateral GFP+ neurons in nIII, yielding 160 IR/MR motoneurons across 17 larvae. The maximum number hit in single larva was 41; dorsoventral distribution remained similar if this larva was removed from consideration, but coverage of dorsal medial nIII (z10-z12) was more sparse. This distribution overlaps minimally with the ventral location of SR and IO motoneurons, supporting our hypothesis that motoneuron somata are orga-nized within nIII along the dorsoventral axis.

### Development of ocular motoneuron organization in nIII proceeds along a dorsoventral order

To determine if the dorsoventral organization that we observed in nIII was related to the time of terminal differentiation, we localized motoneurons born before a series of five time points (22, 29, 36, 43, and 50 hpf). “Birthdating” was accomplished with a photolabile fluorescent protein in a second transgenic line, Tg(huC:Kaede), using a validated technique, Birth-dating Analysis by photoconverted fluorescent Protein Tracing In vivo, with Subpopulation Markers (BAPTISM), detailed in Methods and Figure 4 (Caron et al., 2008). Timepoints spanned most of nIII/nIV nucleogenesis (Higashijima et al., 2000; Clark et al., 2013), allowing us to localize the somata of the majority of neurons in nIII/nIV with respect to their final mitotic division. We used the Tg(isl1:GFP) marker to define nIII/nIV motoneurons, and thus could only localize SR, IR, MR, and SO, but not IO, motoneurons.

We mapped ocular motoneurons born at each time point into the bounds of nIII/nIV (Figure 8). 25-33% of the Tg(huC:Kaede)+ dorsal nIII population evaluated at 5 dpf was born by 22 hpf. Nearly all of dorsal nIII was born by 29 hpf. Almost no ventral nIII motoneurons were born until after 36 hpf; ventral nIII motoneuron birth appeared complete by 50 hpf. We first saw post-mitotic motoneurons in nIV at the 36 hpf conversion, and nIV motoneuron birth also appeared complete by 50 hpf. We conclude that the dorsoventral organization of nIII reflects the birth order of motoneurons.

**Figure 8:**
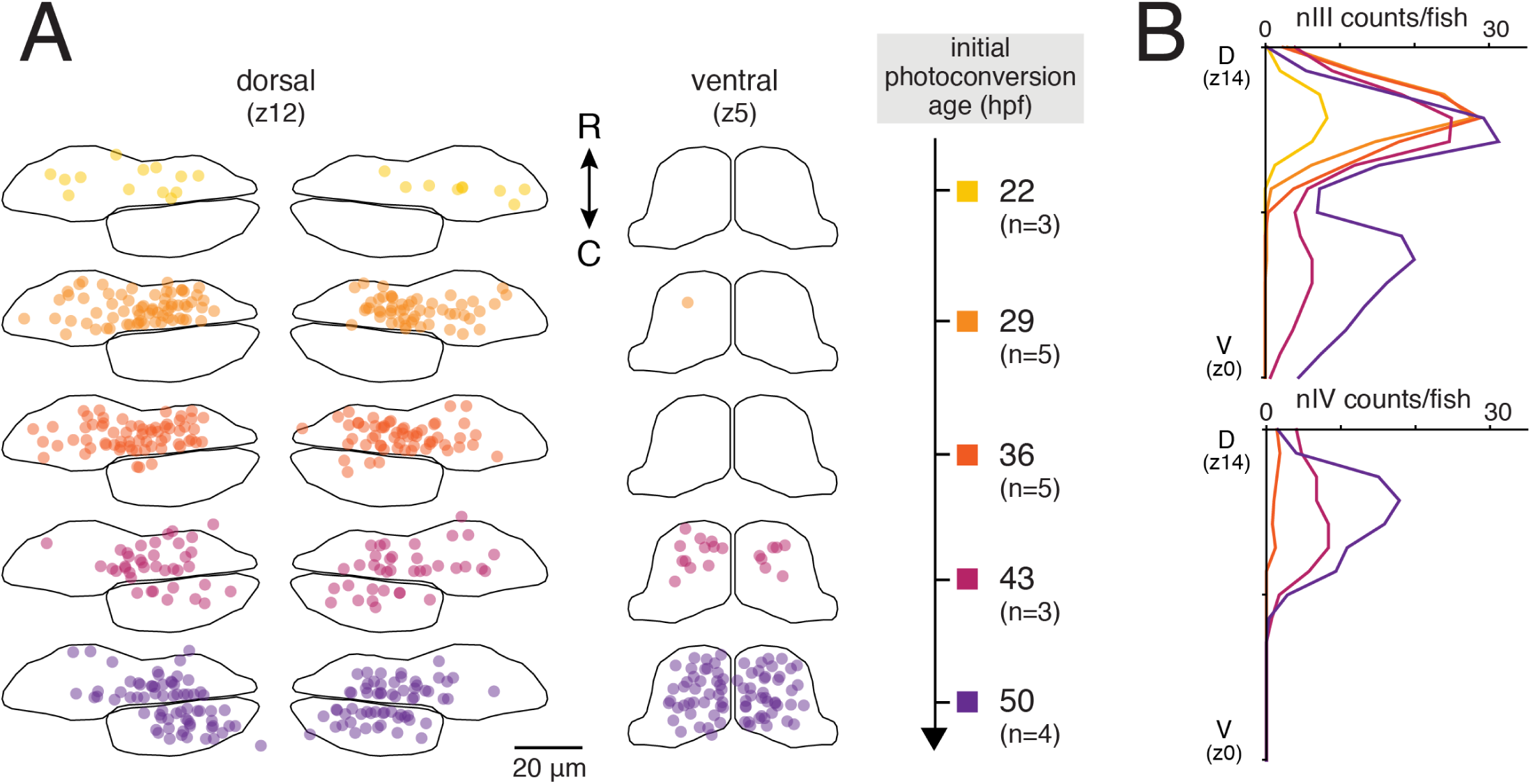
Motoneurons in dorsal nIII are born before those in ventral nIII and nIV. A) Location of motoneurons born by five initial conversion time points, from 22 through 50 hpf, in dorsal nIII/nIV (plane z12) and ventral nIII (plane z5) at 5 dpf. B) Average number of motoneurons born by each time point across nIII and nIV.

## Discussion

This study finds that ocular motoneurons in cranial nuclei III and IV of the larval zebrafish are spatially organized, in a manner consistent with their birth order. We localized ocular motor pools within nIII and nIV of larval zebrafish, assigning identities and relative positions to retro-orbitally dye-filled somata using the characteristic patterns of extraocular muscle innervation and of Tg(isl1:GFP) expression in these nuclei. Our data show a spatial organization to nIII pools in the 5-7 dpf zebrafish: IR/MR motoneurons are predominantly found in dorsal nIII, while IO and SR motoneurons are predominantly found in ventral nIII. Since SR and nIV SO organization does not change between 5-7 and 14 dpf, pool location is likely stable throughout early larval development. By birthdating neurons in nIII/nIV, we have discovered a complementary dorsoventral division in timing of neurogenesis. As both motoneuron identity and birthdate comparably delineate nIII/nIV, the time of terminal differentiation could account for the dorsoventral organization of nIII subpopulations. We propose that birth order plays a preeminent role in determining ocular motoneuron fate, such that dorsal IR and MR motoneurons become post-mitotic earlier than ventral SR motoneurons and SO motoneurons in nIV. Here we consider the limitations of our techniques, a general model for spatiotemporal development of ocular motoneurons, and experiments to elucidate the mechanisms underlying such spatiotemporal organization.

### Limits of the retro-orbital and BAPTISM techniques

Because the larval zebrafish extraocular muscles are too small to restrict dye placement, we classified motoneurons by whether their somata were ipsilateral or contralateral to the targeted eye, and on their overlap with Tg(isl1:GFP) expression. Consequentially, we would incorrectly label any ipsilaterally-innervating SO or SR motoneurons or contralaterally-innervating IO, IR or MR motoneurons. Several earlier studies have reported infrequent ipsilateral innervation of SO or SR, by <5% of the total motoneuron pool in each case (Graf and Baker, 1985; Miyazaki, 1985; Evinger et al., 1987; Sonntag and Fritzsch, 1987; Sun and May, 1993; Murphy et al., 1986; Luiten and de Vlieger, 1978), and one study reported rare contralateral IR motoneurons (El Hassni et al., 2000). The low numbers seen in other organisms lead us to believe that few, if any, motoneurons were incorrectly labeled in our study. We conclude that the spatial organization we observe is likely robust to misclassification errors.

Our protocol does not distinguish IR from MR motoneurons, which may be spatially segregated. Selectively severing only the nasal part of the MR nerve, far from the IR/MR muscle bifurcation, could improve the odds of labeling MR motoneurons. This might allow us to discern a spatial organization to the GFP+ ipsilateral motoneuron population, with the caveat that MR would be incompletely labeled. IR and MR motoneurons do occupy distinct regions of nIII in other species, with varying degrees of overlap. For instance, in other teleosts (goldfish and carp), MR motoneurons have a wider dorsoventral distribution than mainly dorsal IR motoneurons; and MR motoneurons are found clustered ventral to IR motoneurons in flatfish (Graf and McGurk, 1985; Graf and Baker, 1985; Luiten and de Vlieger, 1978). Across vertebrates, IR is usually rostral to MR (Evinger, 1988); and in zebrafish, dorsal nIII may be divisible along the rostrocaudal axis at 50hpf (Clark et al., 2013). We predict, by homology, that zebrafish IR motoneurons may be located rostrally and/or dorsally to MR motoneurons.

The BAPTISM procedure relies on a threshold to separate red Kaede fluorescence indicative of post-mitotic neurons from the basal background level of spontaneous photoconversion. We chose conservative background red Kaede thresholds, which might exclude motoneurons that had only recently exited the cell cycle upon initial photoconversion, whose above-background red Kaede signal would still be faint. Similarly, we may also have missed a cluster of 10-15 post-mitotic motoneurons in ventro-caudal nIII, or 5-8% of total nIII GFP+ cells, that systematically expressed low basal levels of Tg(huC:Kaede). Even accounting for these potential underestimates, our finding of patterned dorsal-to-ventral birth order obtains. Finally, IO motoneurons are not labeled by Tg(isl1:GFP), and so are excluded from our birthdating dataset. We began to see postmitotic GFP-cells around the edges of ventral nIII at the 36 hpf conversion timepoint, but cannot positively identify these as nIII motoneurons. We conclude that the temporal order we observe is likely robust to technical constraints of the BAPTISM method.

### How many motoneurons innervate each extraocular muscle?

Reports from other teleosts vary in the relative numbers accorded to each motor pool (Graf and McGurk, 1985; Luiten and de Vlieger, 1978). While we cannot specify the precise number of neurons that make up a particular motoneuron pool, we propose the maximum number filled in any one larva as an estimate. Based on our Tg(isl1:GFP) counts, nIV is composed of 47±8 SO neurons; the maximum number of SO neurons filled in any one larva was 43, supporting our approach. Within nIII, the maximum number for SR was similar at 41; this would comprise close to the expected one-third (35, total n=104±20) of nIII motoneurons in Tg(isl1:GFP). For IO motoneurons the maximum was only 13, a likely underestimate reflecting the challenge of calling a soma GFP-in the presence of dense GFP+ nIII neuropil and/or signal from surrounding GFP+ cells. Our best estimate for combined IR/MR pool numbers is the total number of GFP+ motoneurons in the dorsal half of nIII, or 5514 per side. This is less than the expected (70) two-thirds o± FP+ neurons in nIII and far less than would be expected relative to SO or SR numbers, given an even subpopulation distribution. The discrepancy between observed and expected numbers could reflect stochastic expression in the Tg(isl1:GFP) line or an unequal number of motoneurons innervating each extraocular muscle. As we never observed a filled motoneuron on the contralateral side that was not GFP+ (i.e. SO and SR motoneurons), Tg(isl1:GFP) expression likely reflects the full complement of motoneurons. Our data is instead consistent with a slight superior/inferior asymmetry, where superior muscles SO and SR may have larger motoneuron pools. At rest, larval zebrafish maintain constant tension in the superior muscles, enabling a slight upward gaze deviation (Bianco et al., 2012), potentially necessitating such an asymmetry.

### A model for the spatiotemporal development of nIII/nIV

Our findings largely support and extend earlier work that defined the timeline for ocular motoneuron axon projection and extraocular muscle innervation in larval zebrafish (compared in Figure 9) (Clark et al., 2013). Our data agree with their observation of neurogenesis in nIII preceding nIV, consistent with the order of *isl1* expression in chick (Varela-Echavarria et al., 1996) but in contrast to their concurrent development in mammals (Altman and Bayer, 1981; Shaw and Alley, 1981). The oculomotor nerve begins projection at 30 hpf, before birth of SR motoneurons has begun. The projecting nerve therefore contains only IR/MR and possibly IO, axons, nIII nucleogenesis is mostly complete by 48 hpf, when the axon front pauses at the edge of the orbit. Because most SR motoneurons are not post-mitotic until after 43 hpf, their axons are likely delayed in reaching this location. Indeed, axon extension toward the IR IO, and MR muscles resumes at 54 hpf, but Clark et al. do not observe a branch innervating the SR muscle until *66* hpf. This may reflect IO differentiation before SRin the zebrafish, in agreement with the order seen in rabbit (Shaw and Alley, 1981).

**Figure 9:**
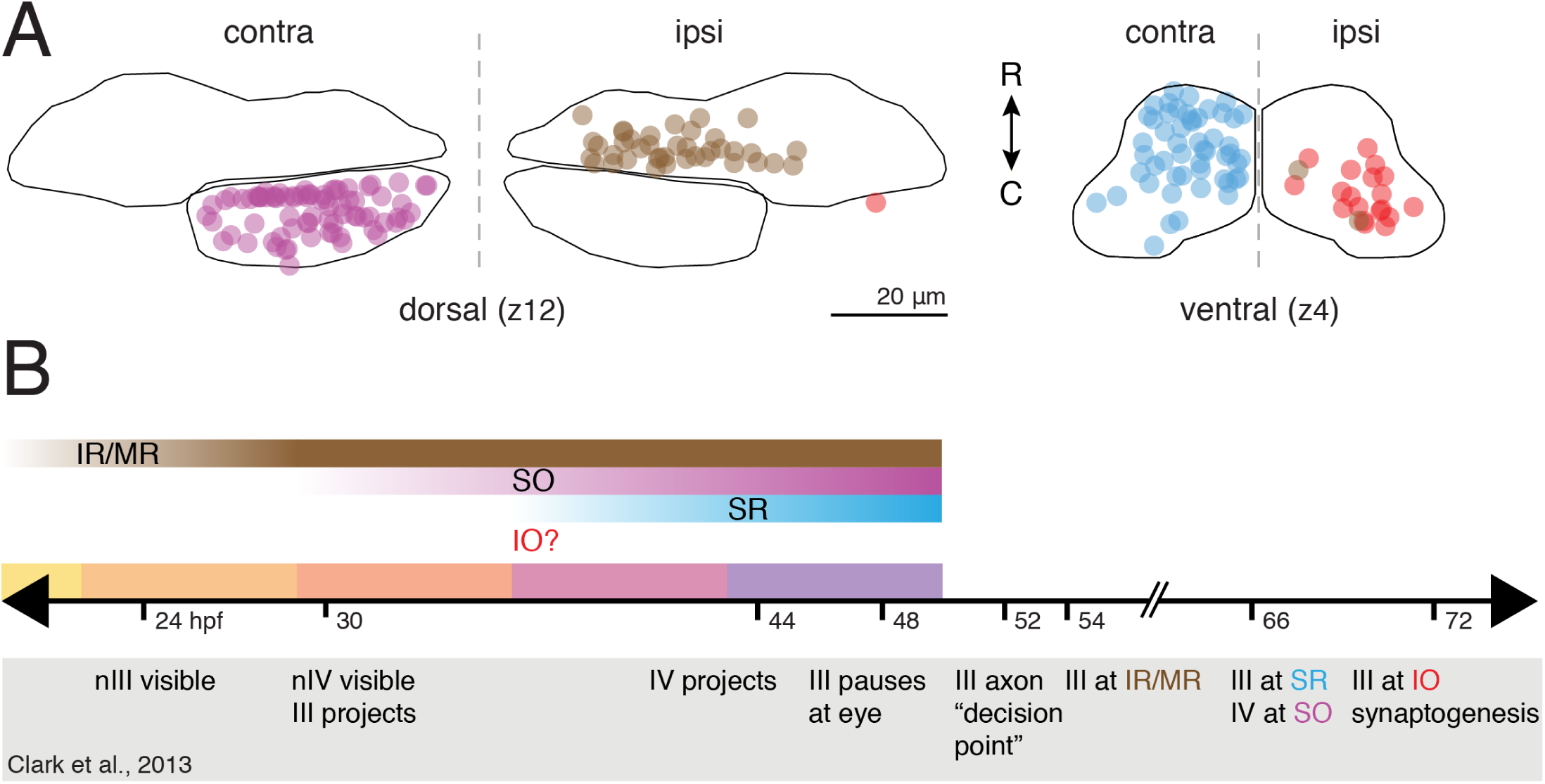
Spatiotemporal organization of ocular motoneurons in nIII and nIV. A) Aggregate figure showing relative locations of nIII/nIV motoneurons at two exemplar planes; data from Figs. 6 & 7. B) Comparison of motoneuron subpopulation birthdates (gradient bars) with development landmarks (Clark et al., 2013) from live imaging of ocular motoneuron projections (timeline markers; events inside grey box). Solid bars: periods between BAPTISM initial photoconversion time points.

Our data speak to two prior hypotheses about the developmental organization of ocular motoneurons. One proposition was that a shared caudal-to-rostral sequence of ocular motoneuron and extraocular muscle development allows progressive innervation of earliest-born muscles by earliest-born ocular motoneurons (Evinger, 1988). This hypothesis does not match the true order of events, since the IR and MR muscles, which are the latest to differentiate (Noden et al., 1999) and still a paired anlagen when invaded by motoneuron axons (Clark et al., 2013), are innervated by the earliest motoneurons to differentiate. In contrast, nIV motoneurons innervating the earlier-differentiating SO muscle are born only after IR/MR motoneuron birth is complete. Nor do our results agree with a random innervation model of nIII development (Glover, 2003). Such a model predicts no correlation between motoneuron birthdate and muscle target, in contrast to the dorsal-to-ventral progression we observe. Other factors thus are likely to define the organization of nIII/nIV, at least in larval zebrafish.

The dorsoventral organization of nIII in larval zebrafish is similar to that seen in avians and other teleosts (Evinger, 1988); embryonic avian nIIIs feature particularly well-delineated dorsolateral IR, dorsomedial MR, and ventral IO (lateral) and SR (medial) regions. The timing of nIII development we observe likewise agrees with previous studies: Hasan et al., (2010) found a dorsal-to-ventral gradient of neurogenesis in the chick oculomotor complex (which includes Edinger-Westphal motoneurons). Since avian and zebrafish nIII dorsoventral spatial organizations are comparable, the dorsoventral temporal gradient in chick implies birth of IR/MR before IO/SR motoneurons, as we report here. Shaw and Alley (1981) found an IR-MR-IO-SR progression in the birth order of rabbit nIII subpopulations. In rabbits and other mammals, the dorsoventral spatial organization is inverted relative to avians and teleosts, with IR and MR populations generally ventral to IO and SR populations. The temporal order of motoneuron birth thus appears to be conserved, while migration pattern/eventual location within the nucleus may not be. We propose that our finding in the larval zebrafish of motoneuron pools organized according to birth-date is likely general, even when the final spatial layout is different.

### Potential mechanisms and consequences of spatiotemporal development

Spatiotemporal determination of identity is a fundamental theme of neurogenesis (Moore and Livesey, 2015). One possible mechanism for temporally ordered development is to ensure that motoneuron precursors sit in particular spatial locations within nIII, where external cues then specify identities of each subpopulation (Shaw and Alley, 1981). Experiments in which oculomotor complexes were ectopically induced laterally and rostrally in the chick midbrain, with normal dorsoventral positioning of visceral vis-a-vis somatic MNs, suggest that major spatial cues working on the developing nIII lie along the dorsoventral axis (Hasan et al., 2010). Other extrinsic contributions might resemble the negative feedback cues onto retinal cell progenitors that inhibit prolonged production of early-born retinal ganglion and amacrine neurons (Belliveau and Cepko, 1999; Waid and McLoon, 1998; Poggi et al., 2005), as retinoic acid produced by early-born spinal cord motoneurons acts on progenitors as a cue for late-born motoneuron generation (Sockanathan and Jessell, 1998). However, cell-autonomous intrinsic factors such as the “temporal series” of transcription factors inherited by the offspring of *Drosophila* neuroblasts (Isshiki et al., 2001; Jacob et al., 2008) may also play a role. Transplantation of GFP+ neurons from Tg(isl1:GFP) donor embryos into particular spatial locations in the dorsal or ventral midbrain would allow us to differentiate intrinsic and extrinsic factors, as well as to discover the precise timing of ocular motoneuron commitment. Similar experiments performed in cortex (Frantz and McConnell, 1996; McConnell and Kaznowski, 1991) and spinal cord (Appel et al., 1995) have defined time windows within which progenitors’ competence to respond to external cues shifts or narrows, representing a convergence of intrinsic and extrinsic cues.

As the final target for all oculomotor circuits, multiple upstream inputs must find partners within particular ocular motoneuron pools. Our data support a model in which, by the time motoneurons occupy their final positions within nIII, they possess sufficient information to choose a target extraocular muscle. Our study thus supports molecular profiling of developing motoneurons at this time, specifically a comparison of dorsal and ventral GFP+ neurons from Tg(isl1:GFP) at later stages of nucleogenesis. We hypothesize that each subpopulation will express a unique complement of terminal effectors (Hobert, 2016) that allow it to respond appropriately to cues arising from potential muscle targets responsible for motoneuron matching. Similarly, our model suggests that motoneurons may be competent to signal upstream partners, even before synaptogenesis onto target extraocular muscles (72 hpf). Since the motoneurons occupy fairly stereotyped regions, neurons projecting to the ocular motor nuclei might themselves rely on spatial cues to aid targeting, a strategy employed by sensory afferents projecting to the expected locations of motor pools in the spinal cord (Sürmeli et al., 2011). For instance, vestibular projection neurons mediating the “nose-down/eyes-up” response of the vestibulo-ocular reflex (joint activation of IO and SR motoneurons/muscles) would be able to target ventral nIII with little chance of incorrect innervation (Bianco et al., 2012). Spatial segregation is likely to complement rather than replace more traditional molecular recognition cues, due to imperfect segregation of motoneuron pools within nIII.

Together, transplantation and molecular profiling experiments can uncover candidate mechanisms and factors downstream of *isl1* expression that define ocular motoneuron subtypes. Understanding the contributions of temporal and spatial factors in nIII motoneuron identification should clarify how they find, and are found by, partners during circuit construction. Our data showing common dorsoventral organization between motoneuron identity and birth order provides crucial spatial and temporal guidance for this set of experiments. We propose that the larval zebrafish ocular motor system is a powerful model to uncover the molecular basis for ocular motoneuron organization, and by extension, the behaviors it subserves.

## Acknowledgments

Research was supported by the National Institute on Deafness and Communication Disorders of the National Institutes of Health under award number 5R00DC012775. We thank Fernando Fuentes, Katherine Harmon, Eva Lancaster, and Belinda Sun for their help in caring for zebrafish lines used in this study; and Bob Baker, Jeremy Dasen, Holger Knaut, Katherine Nagel, David Ehrlich, and Katherine Harmon for comments on the manuscript.

## Conflict of Interest Statement

The authors declare no conflicts of interest.

## Author Contributions

Conceptualization: MG and DS. Methodology: MG and DS. Investigation: MG, AP, and KD. Validation: MG and AP. Visualization: MG. Writing: MG and DS. Funding Acquisition: DS. Supervision: DS.

